# Intra-voxel incoherent motion magnetic resonance imaging of the living human fetus: the technique and within-subject reproducibility

**DOI:** 10.1101/180844

**Authors:** András Jakab, Ruth Tuura, Raimund Kottke, Christian Kellenberger, Ianina Scheer

## Abstract

Our purpose was to test the within-subject repeatability of the perfusion fraction, diffusion coefficient and pseudo diffusion coefficient measurements in various fetal organs and in the placenta based on the intra-voxel incoherent motion imaging principle. *In utero* diffusion-weighted magnetic resonance imaging was performed on 1.5T and 3.0T clinical scanners with b-factors ranging from 0 to 900 s/mm2 in 16 steps and a tetrahedral diffusion-weighting encoding scheme. Data from 16 pregnant women (maternal age: 34 ± 4.9 years, range: 24.6 −40.8) were included in this pilot study. For 15 cases, IVIM was repeated (maternal age: 33.7 ± 5.2 years, range: 24.6 – 40.8). A bi-exponential model was fitted on the volume-averaged diffusion values and the perfusion fraction (f), diffusion coefficient (d) and pseudo diffusion coefficient (D*) were calculated. Within-subject repeatability was given as the test-retest variability of the IVIM parameters in the fetal frontal cortex, frontal white matter, cerebellum, lungs, kidneys, liver and in the placenta. An in-house developed image processing script was utilized to remove the image frames with excessive motion, and to perform motion correction by using non-linear freeform deformations. For the fetal lungs, liver and the placenta, within-subject variability ranged from 14.4% to 20.4% for f, 12.2% to 14.1% for d and 16.8% to 25.3% for D*. The diffusion coefficients of the investigated brain regions were moderately to highly reproducible (4.8% to 15.2%), however, f and D* showed inferior reproducibility compared to corresponding measures derived for the lungs, liver and placenta. The IVIM-based parameters of the fetal kidney were revealed to be highly variable across scans. Our results indicate that in utero intra-voxel incoherent motion magnetic resonance imaging potentially provides a novel method for examining microvascular perfusion and diffusion in the developing human fetus. The reproducible quantification of the perfusion and diffusion parameters depend greatly upon data quality, fetal and maternal movements, and image post processing to detect and remove corrupted data before calculating the IVIM model.

## INTRODUCTION

Early pioneers of nuclear magnetic resonance (NMR) found that the MR signal can be sensitized to the displacement of water molecules caused by the thermally driven self-diffusion process (Stejskal, 1965). This observation led to the development of magnetic resonance imaging (MRI) sequences probing diffusion-driven displacements in living biological tissues (Merboldt et al., 1991), such as the diffusion-weighted imaging (dMRI or DWI) method, which sensitizes MR images to water diffusion with pulsed magnetic gradients (Le Bihan et al., 1986). During DWI, the amount of diffusion-weighting is quantified with the b-factor which describes the degree to which diffusion processes contribute to the MR signal. By acquiring diffusion weighted images (DWI) with one non-zero b-factor and (at least) three diffusion gradient directions, it is possible to quantify the apparent diffusion coefficient (ADC), a measurement that has important implications for the diagnosis and assessment of neurological disorders (Schlaug et al., 1997).

Biological tissues, however, exhibit more complex diffusion characteristics, which cannot be fully described using a single, directionally uniform diffusion coefficient (Jensen et al., 2005; Niendorf et al., 1996). To resolve the directional non-uniformity of diffusion in tissues such as the white matter fibers in the brain, diffusion tensor imaging can be used to probe diffusion using pulsed magnetic gradients of at least six varying spatial directions. Tensors or other more complex mathematical objects can provide models for the diffusion characteristics observed during such measurements (Basser et al., 1994). However, while the tensor model can characterize the directional variability of the diffusion signal, the presence of multiple anatomical compartments, complex barriers or other structures affecting the microscopic displacement of proton spins can result in multiple diffusion coefficients coexisting in the elementary imaging units (voxels), such that the mono-exponential model of diffusion may not unambiguously represent the underlying physiological phenomena.

The **intra-voxel incoherent motion (IVIM**) concept describes micro-scale translational movements within imaging voxels with a bi-exponential model. While thermally-driven Brownian motion results in relatively low apparent diffusion coefficients in human tissues, water protons in biological tissues can also undergo similar displacements as a result of their perfusion-driven flow across the microvascular network. The latter process causes significantly larger displacements per unit time and an order of magnitude larger diffusion coefficient than the “true” (thermal) diffusion, forming the basis of the bi-exponential theory underlying IVIM. The possible applicability of the IVIM concept in diagnostic imaging was initially suggested in 1989 by Le Bihan et al. (Le Bihan et al., 1988; Le Bihan and Turner, 1992). More recently, faster MRI sequences have paved the way for clinical applications of IVIM (Federau et al., 2012; Le Bihan, 2012; Lemke et al., 2010).

IVIM relies on the assumption that the fast-moving component arises from blood flowing across the vascular bed in such a way that mimics a random walk (hence the word “incoherent”). To meet this criterion, the vascular spaces must have uniform orientation distribution within voxels. Studies based on IVIM report that the separation of diffusion and perfusion allows more accurate estimation of tissue diffusivity (quantified as the ***slow diffusion coefficient, or real diffusion, indicated as d or ADCiow)*** in organs that are intrinsically highly perfused, such as the liver, kidney or placenta. The diffusion coefficient corresponding to the fast component is commonly regarded as the ***pseudo-diffusion coefficient or fast diffusion coefficient; D* or ADCfast.*** A third important parameter, the ***perfusion fraction (f)***, describes the fraction of incoherent signal arising from the vascular compartment in each voxel. The relationship between the signal intensity change caused by both diffusion processes and diffusion coefficients is commonly formulated as:

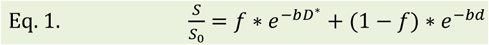

*where S is the measured signal intensity, SO is the signal intensity without diffusion-weighting, d is the diffusion coefficient, D* is the pseudo-diffusion coefficient, f is the perfusion fraction and b is the b-factor.*

The parameter *f* is more likely to represent the relative amount of blood flowing through the vascular bed rather than the flow velocity itself, and Le Bihan et al. established a direct relationship between cerebral blood volume and perfusion fraction using the following equation (Le Bihan and Turner, 1992):

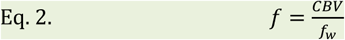

*where f is the perfusion fraction, CBV is the cerebral blood volume and f****w*** *is the MRI visible water content fraction (an empirical constant).*

Preliminary studies demonstrated the use of the IVIM imaging in pregnancy (Moore et al., 2000) by characterizing the effect of pathological conditions on the IVIM parameters, such as the alteration of the perfusion fraction associated with intrauterine growth retardation (Alison et al., 2013; Derwig et al., 2013; Siauve et al., 2015). Similarly, arterial spin labelling by means of flow sensitive alternating inversion recovery (ASL-FAIR) has been shown to offer a method for evaluating the transit of blood across the placenta (Derwig et al., 2013).

The successful adaptation of the IVIM technique to the prenatal imaging setting would have important implications for clinical decision-making. For example, the perfusion in the microvascular compartment of the developing fetal lungs, brain, kidneys or other organs may serve as an indicator of vascular development and organ viability. However, the acquisition of high quality imaging data with DWI or other echo-planar imaging-based sequences *in utero* is extremely challenging due to subject motion and the complex anatomical and biochemical environment. Here we present a novel work-flow for the acquisition and analysis of fetal IVIM images. Our purpose was to test the within-subject repeatability of the perfusion fraction, diffusion coefficient and pseudo diffusion coefficient in various fetal organs and in the placenta. While previous prenatal studies focused on fetal DWI with relatively high b-factors (e.g. 500 to 800 s/mm^2^) (Balassy et al., 2008; Lee et al., 2009; Moore et al., 2001; Savelli et al., 2007; Schneider et al., 2009), we aimed to implement the IVIM principle using diffusion weighting with additional lower b-factors, which may allow the separation of real diffusion and perfusion effects, and thus improve the specificity of the detection of pathological changes in microvascular perfusion of the fetal organs *in utero.*

## MATERIALS AND METHODS

### Patients

MRI data from 16 pregnant women (maternal age: 34 ± 4.9 years, range: 24.6 - 40.8) were included in this pilot study. For 15 cases, IVIM was repeated (maternal age: 33.7 ± 5.2 years, range: 24.6 - 40.8). In two cases, fetal MRI was performed at two time points during gestation with 2 and 2.5 and weeks between scans-these two measurements of the same cases were treated as independent samples, resulting in 17 IVIM data sets in total for the reproducibility analysis. The gestational age of the fetuses was 26.3 ± 3.7 (21-36) weeks. Fetal MRI was clinically indicated in all cases to rule out or confirm suspected pathologies detected during prenatal screening by ultrasonography. The clinical indication for MRI was isolated mild ventriculomegaly in 5 cases, myelomeningocele in 8 cases, and 1 case each of sacrococcygeal teratoma and congenital bronchial atresia.

### Ethical statement

The mothers gave written informed consent before the MRI examination, and the Ethical Commission of Canton Zürich approved the study (EK Number: 2017-00167).

### MR imaging

Fetal MRI was performed on two different clinical MRI systems as part of the routine clinical assessment (1.5 GE Discovery MR450: 12 datasets, 3.0T GE Discovery MR750: 6 datasets). The assignment of the cases to an individual scanner was not controlled in the current study, and was based on the availability of free scanner time. IVIM data were collected from January 2016 until March 2017 at the University Children’s Hospital Zürich. Pregnant women were examined in the supine position, feet first, and no contrast agents or sedatives were administered. In order to obtain optimal MR signal for the fetal head and body within the same session, the coil was readjusted to the position of the fetal structures investigated.

For each fetus in this study the IVIM imaging sequence was repeated twice with identical settings. The sequence relied on a DWI sequence that was optimized for fetal imaging, and was modified to accommodate more b-factors within a feasible imaging time. Slices were positioned in axial plane relative to the fetal brainstem (in case of brain imaging) or in coronal plane relative to the fetal body (when imaging fetal organs and the placenta).

A dual spin-echo echo-planar sequence was used with TE/TR: 2200/75 ms, acquisition matrix: 80*100, voxel size: 2*2 mm, slice thickness: 3-4 mm, slice gap: 0.5 mm, number of slices: 8-14, NEX: 1. The “tetra” (tetrahedral) diffusion-weighting orientation scheme was used, which utilizes four different combinations of X,Y and Z diffusion gradients (Conturo et al., 1996). B-factor values were increased in 16 steps and one B0 image was acquired (b-factors: 0, 10, 20, 30, 40, 60, 80, 100, 150, 200, 300, 400, 500, 600, 700, 800, 900 s/mm^2^). This scheme resulted in 64 diffusion-weighted images and one B0 image in each IVIM image series. The actual imaging time depended on the number of slices, which was adjusted to the size of the fetus, and focus of the investigation, or specifically whether the brain (8 to 12 slices) or the whole fetal body and placenta (10 to 15 slices) were the most important organs for clinical decision making. Imaging time per IVIM acquisition ranged from 1:40 to 3:20 minutes.

### IVIM image processing

The processing of fetal IVIM images was carried out using an in-house developed script written in BASH language for Linux. It utilized image processing algorithms from the FSL (Jenkinson et al., 2012), C3D and NIFTIREG software packages for image registration and re-sampling. The image analysis script is available as **digital supplement** to this manuscript. First, the raw IVIM data were viewed using the *fslview* command of the FSL software, and the image frames with the most excessive subject motion were marked and removed from the analysis. This step was followed by a nonlinear, free-form deformation-based registration of image frames with the reg_f3d command in the NIFTIREG tool (Modat et al., 2010), the registration steps of which are illustrated in **Figure 1**. This image registration step used a fine deformation grid with a grid spacing of 6 * 6 * 6 mm for the low b-factor image frames and 12 * 12 * 12 mm grid for the high b-factor frames.

**Figure 1.**
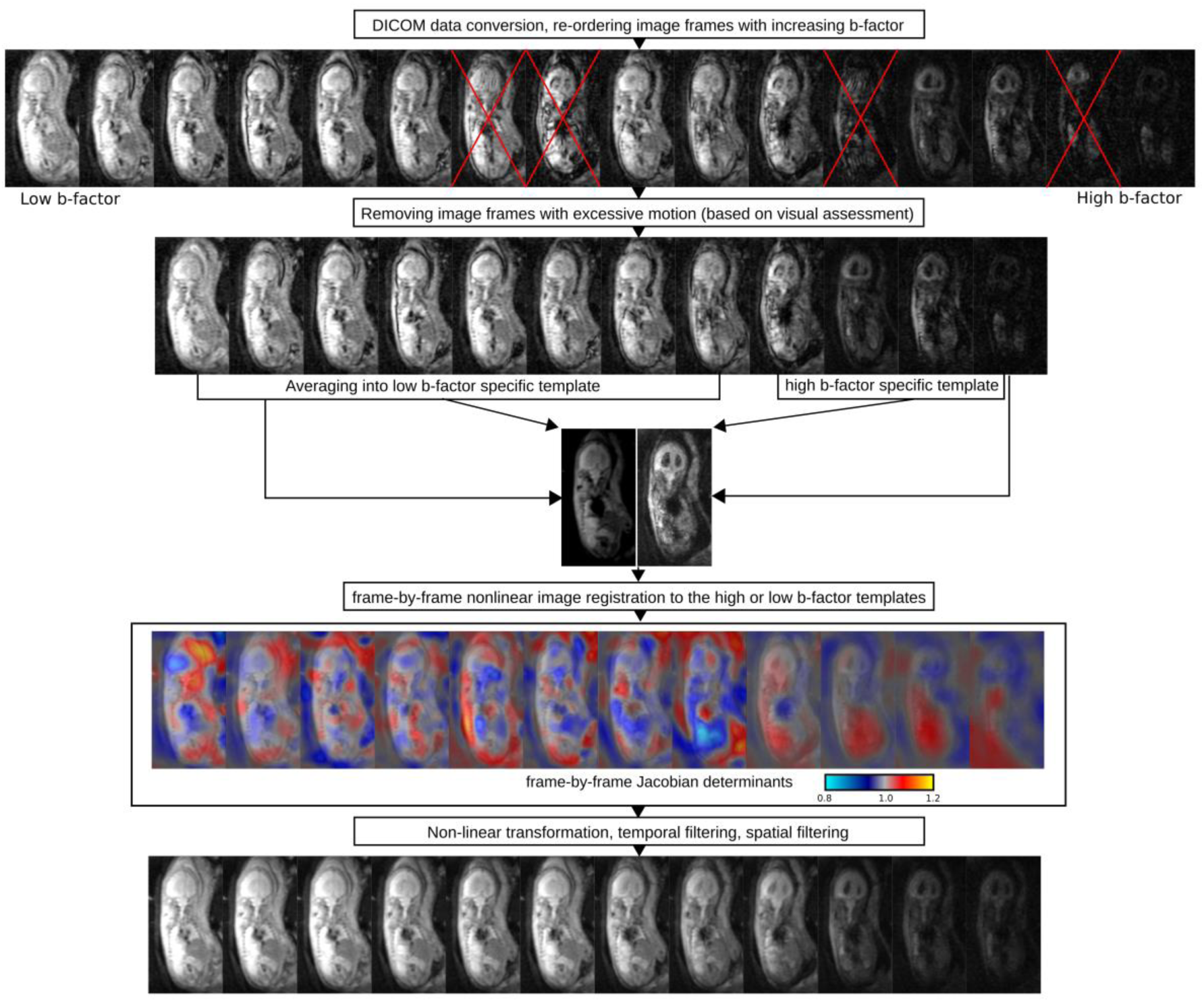
Image processing steps to correct fetal in utero IVIM data sets for subject motion.

### Volume of interest definition

After processing the IVIM data, averaged images for low (b<250 s/mm^2^) and high b-factors (500-900 s/mm^2^) were created. Using the manual segmentation tool in the MITK diffusion toolkit (Wolf et al., 2005), volumes of interest (VOI) were placed over the central parts of the placenta, on the fetal liver, lung parenchyma excluding the hili, kidneys bilaterally, cerebellum and brainstem, frontal or frontoparietal cortical mantle and white matter of the frontal and parietal lobes (**Figure 2**). VOIs were drawn manually by one observer (A.J.). 3D interpolation in the MITK software was then used to smooth the borders of the delineated organ labels. For the kidneys and the placenta, better visual discrimination from surrounding tissues was achieved by delineating the ROIs by viewing the diffusion images with higher b-factors, while for the other structures, we used the diffusion MRI images that were averaged over lower b-factors.

**Figure 2.**
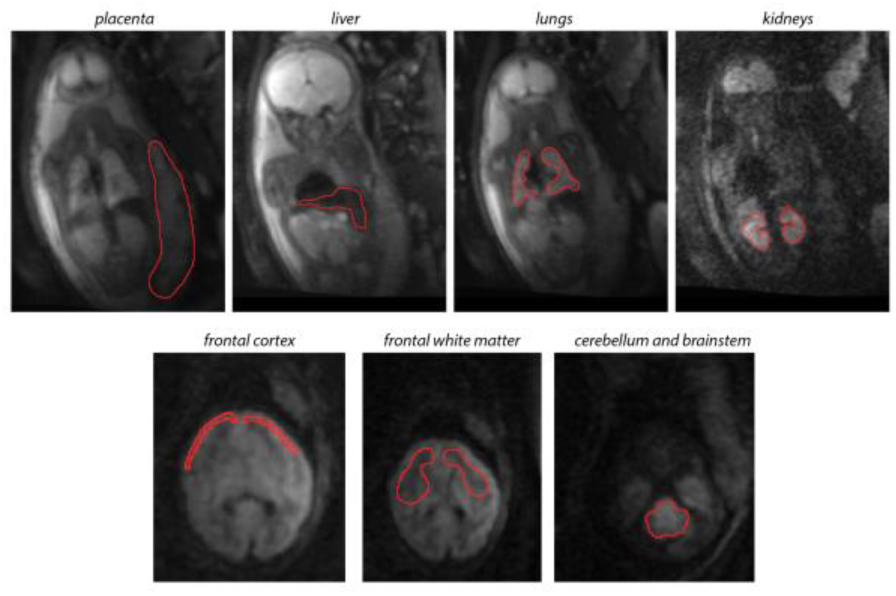
Volume of interest (VOI) delineation of various fetal organs and the placenta. VOIs have been manually delineated to test the within-subject reproducibility of the parameters that are calculated from IVIM data. Red overlay: manual outlines of the organs, background image: coronal or axial diffusion-weighted images.

### IVIM model fitting

The IVIM parameters *f, d* and *D\** (see Introduction) were estimated based on the VOI-averaged signal intensity values to achieve a better signal-to-noise ratio. The analysis of diffusion and perfusion parameters with the IVIM model assumed two compartments with no interactions between the compartments (Federau et al., 2012), and a bi-exponential model (Eq. 1.) was fitted in two steps on the averaged signal intensity over the VOIs. First, the measurements were fitted for b-values larger than 250 s/mm^2^ to estimate the parameter *d* using a mono-exponential term. Then the *f* and *D\** were estimated keeping *d* fixed at the previously fitted value. The IVIM model fitting was carried out with the MITK diffusion toolkit.

### Reproducibility analysis

Reproducibility of the *f, d* and *D\** over the repeated scans were given as the test-retest variability:

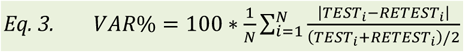

where *N* is the number of subjects *TEST*_*i*_ and *RETEST*_*i*_ are the duplicate measurements for subject *i.*

Next, we tested whether the variability of the IVIM parameters is affected by possible confounds. Multiple, univariate ANOVA tests were carried out with the “General linear model” module in SPSS v22.0 for Windows (Mathworks inc., Nattick, MA, USA). In this analysis, the test-retest difference (that is: 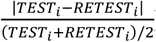) of the *f, d* and *D\** of each organ served as the dependent variable. We evaluated the effect of gestational age, maternal age, scanner field strength and the number of removed image frames on the test-retest difference of each IVIM parameter of the investigated organs. To reveal interactions between the IVIM parameters and the assumed confounds, we report results of the ANOVA tests where the level of significance was p<0.05.

## RESULTS

### IVIM imaging characteristics of fetal normal and pathological tissues and the placenta

**Figure 3/a** demonstrates the typical appearance of the diffusion-weighted fetal images during an IVIM sequence in well-perfused organs: with increasing b-factor, the image intensity decreases exponentially, especially in the higher b-factor range (>250 s/mm^2^, illustrated as red dots in the diagram). In the lower b-factor range (<250 s/mm^2^), the diffusion appears to be faster and shows higher S/S0 ratio than what would be expected by extrapolating the the mono-exponential fitting (black dots and black regression line). The IVIM images in the low b-factor range show predominantly T2 characteristics, and most fetal organs were easy to delineate. With increasing b-factor, only the brain, kidneys, placenta and the muscular layer of the uterus remained distinguishable from the background noise (**Figure 3/a**, second row).

**Figure 3.**
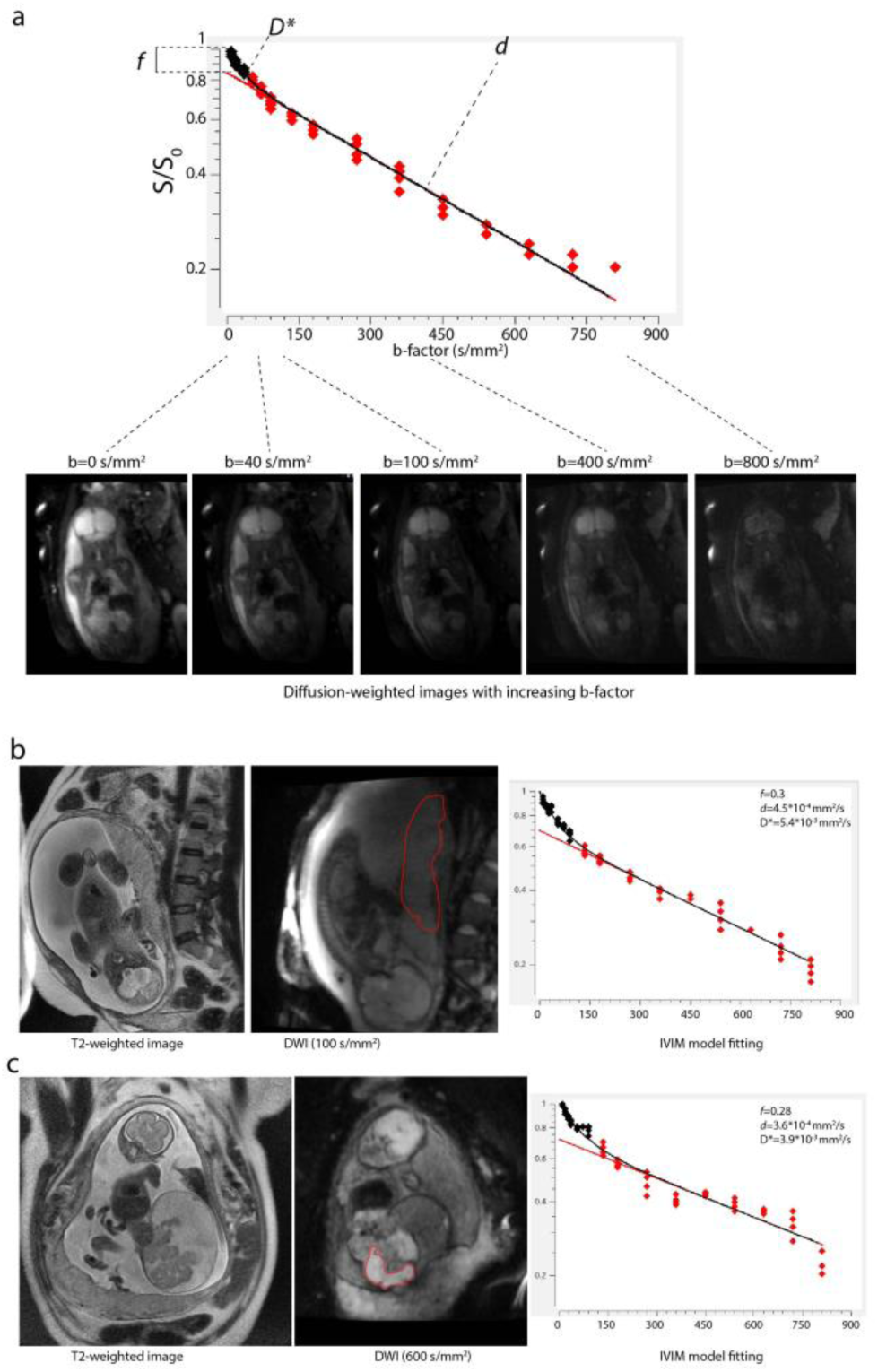
IVIM imaging in utero. ***(a)*** *fitting process of a bi-exponential model on the individual DWI measurements that have been acquired with increasing b-factor*, ***(b)*** *IVIM imaging of the placenta, left image: T2-weighted MRI in coronal plane, middle image: DWI image and placenta delineated, right image: the IVIM signal and estimates of the diffusion coefficient “d”, pseudo-diffusion coefficient “D*” and the microvascular perfusion fraction “f”, based on VOI-averaged values.* ***(c)*** *IVIM imaging of a sacrococcygeal teratoma.*

We found high microvascular perfusion fraction in the fetal liver (f=0.346±0.101) and lungs (f=0.33±0.112). The liver appeared as a homogeneously and highly perfused organ on the IVIM parametric maps. The central (hilar) parts of the lungs displayed higher *f* than their periphery, while the *f* was not as prominently high as in the adjacent heart and great vessels, which typically had *f* values over 0.5. The central parts of the placenta were moderately and homogeneously perfused (population average f=0.28±0.105), with a tendency towards higher perfusion near the the basal layer of the placenta (**Figure 4**, asterix). The kidneys were also moderately perfused (f=0.153±0.09). We found high heterogeneity and putative partial volume artifacts caused by the movement of these organs relative to the imaging plane due to maternal breathing, fetal breathing and fetal trunk movements.

**Figure 4.**
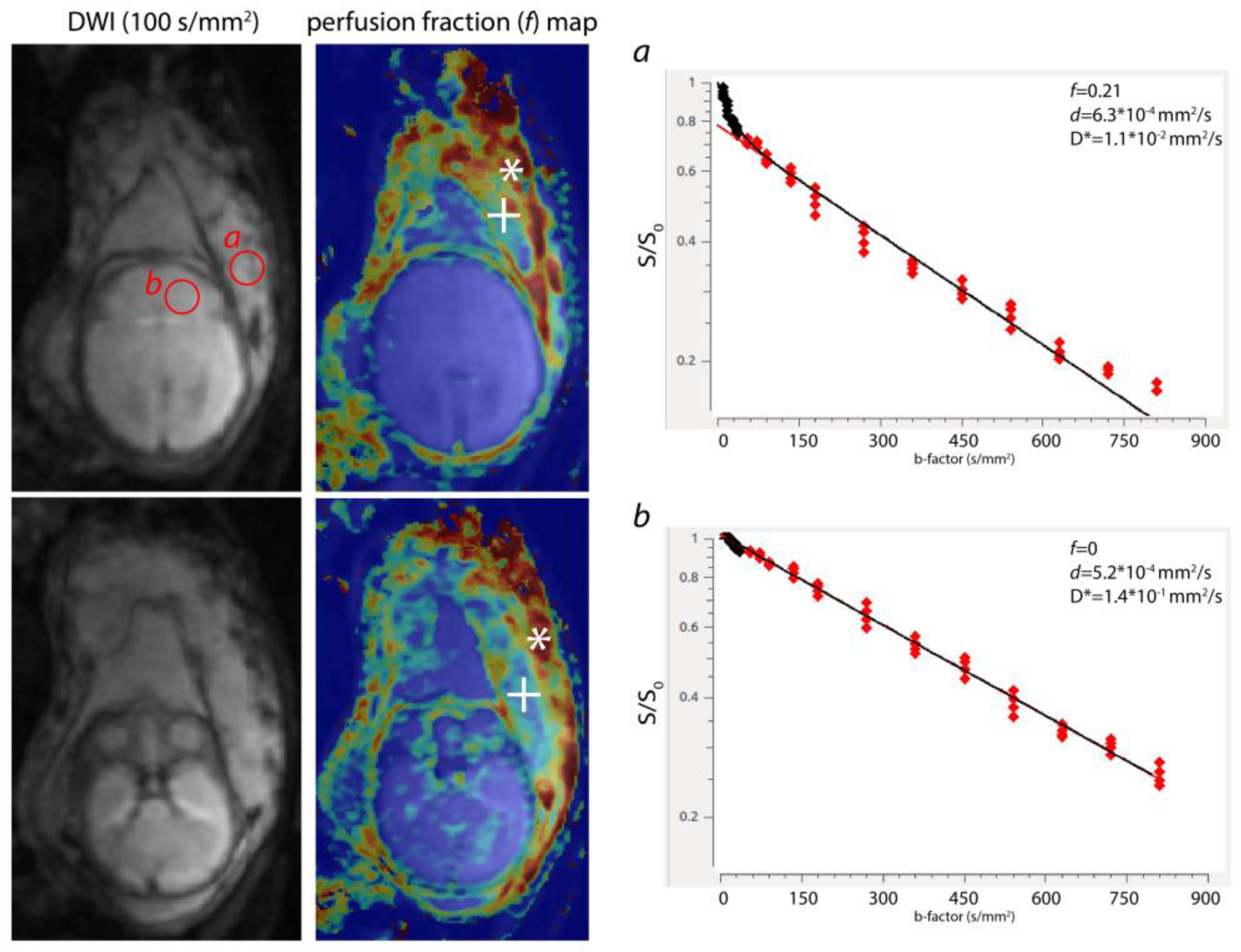
IVIM imaging of the fetal brain. *Images on the left: diffusion-weighted axial images showing the fetal brain, middle images: perfusion fraction (f) maps, right images: bi-exponentialfitting based on the IVIM measurements. Compared to the neighboring placenta (a) and especially its basal plate* (central part of placenta: +, basal plate: *), *we measured very low f or D* values in the fetal brain (IVIM curve:* ***b).***

All three regions of the brain demonstrated low microvascular perfusion (frontal white matter, f=0.087±0.106; frontal cortex, f=0.142 ± 0.133; cerebellum, f=0.135 ± 0.122). Interestingly, in many cases fand *D\** in the fetal brain were estimated to be 0, which is most likely an artefact due to the insufficient data quality for estimating low perfusion values, or *D\** was estimated to be smaller than *d.* Compared to the neighboring central placenta or basal plate (**Figure 4**, cross and asterix, respectively), the fetal brain, especially the white matter, appeared to be almost nonperfused (**Figure 4/b**). We measured higher *f* values in the fetal brain’s frontal cortex, however, this is likely to arise from partial volume effects with the adjacent cerebro-spinal fluid spaces, which inherently display sharp signal decay in DWI experiments due to the bulk movement of proton spins.

### Within-subject reproducibility

Images acquired with the IVIM protocol were prone to three main sources of motion and consequent artifacts: maternal breathing, fetal body movement and physiological movements of fetal internal organs, which can be identified by looking at the raw diffusion-weighted images or observing the outlier points (**Figure 5**, arrows) during the bi-exponential curve fitting procedure.

**Figure 5.**
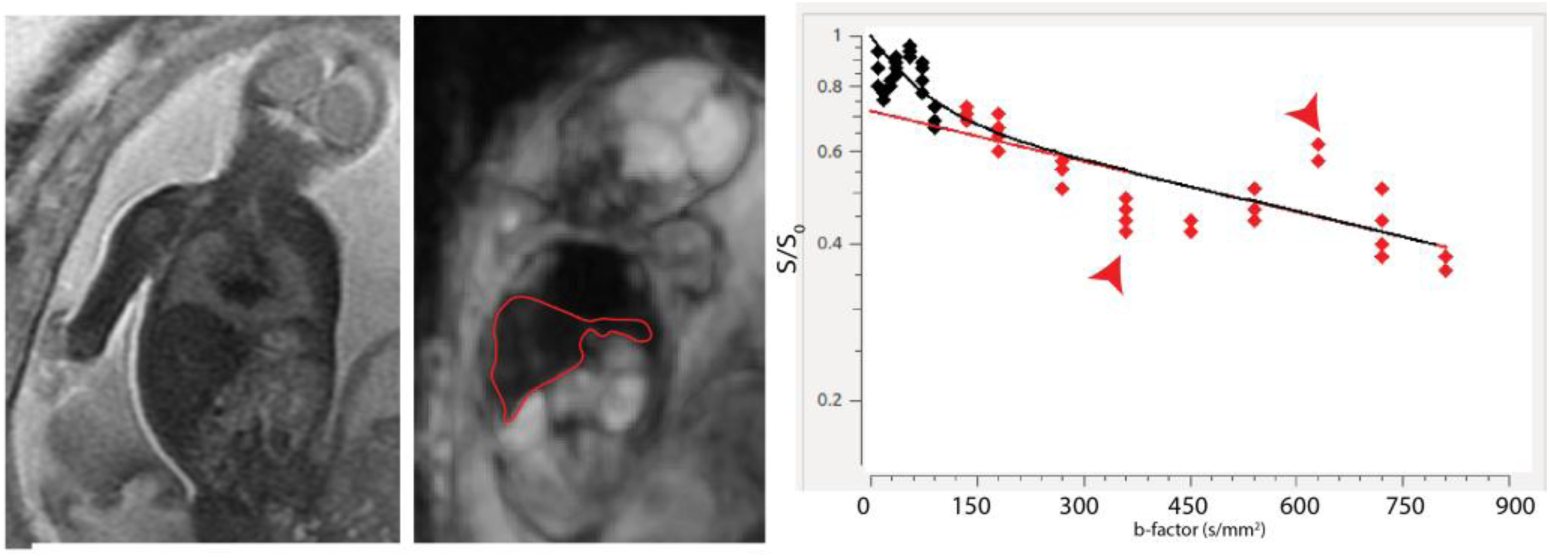
IVIM imaging of a fetus that has moved considerably during the acquisition. *Left image: T2-weighted MRI of the fetus, middle image: diffusion-weigh ted image, righ t image: bi-exponential fitting based on the IVIM measurements. Sudden changes in pose and fetal breathing movements cause particularly large displacements of fetal abdominal organs, such as the liver (red outline, middle image) and increase- or decrease the measured signal intensity (red arrows). To overcome this, frame-by-frame visual inspection may become necessary.*

The effect of large movements of the fetal body was partially mitigated by removing 6.1 ± 7.4 (CI: 0-20.1) image frames with excessive motion before the analysis. Reproducibility of the diffusion coefficient was the highest for all the investigated fetal structures and the placenta (VAR%, frontal cortex: 4.8%, placenta: 12.2%), as estimated by the mono-exponential decay of signal intensities corresponding to image frames with a b-factor >250 s/mm^2^. In contrast, *f* and *D\** values were twice as variable across repeated scans as the *d.* Only three organs, the placenta, fetal liver and lungs were found to show moderately reproducible perfusion fractions and pseudo-diffusion coefficients (Table 2), while fand the *D\** of the investigated brain areas and the kidneys showed poor reproducibility (with a test-retest variability over 25%).

**Table 2.** *Fetal IVIM parameters and their reproducibility across the study population. Data are given as the mean ± standard deviation (SD) and value range, while reproducibility is expressed as test-retest variability (VAR%, Eq. 3.) of duplicate measurements.*

| Structure<br>(number of<br>subjects where<br>structure was<br>repeatedly<br>imaged) | IVIM measurements | | | Within-subject variability of IVIM<br>measurements | | | Subjects<br>where bi-exponential<br>fitting was<br>insufficient<br>($f$ and<br>$D^*=0$;<br>$n$ / %) |
| --- | --- | --- | --- | --- | --- | --- | --- |
| | Perfusion<br>fraction $f$,<br>unitless,<br>mean ± SD<br>(min-max) | Diffusion coefficient<br>$d$, mm<sup>2</sup>/s, mean ±<br>SD, (min-max) | Pseudo-diffusion<br>coefficient $D^*$,<br>mm<sup>2</sup>/s, mean ± SD,<br>(min-max) | Perfusion<br>fraction $f$,<br>VAR% | Diffusion<br>coefficient<br>$d$, VAR% | Pseudo-diffusion<br>coefficient<br>$D^*$, VAR% | |
| Brain, frontal<br>cortex<br>($n=11$) | **0.142** ± 0.133<br>(0 – 0.54)* | **4.7\*10<sup>-4</sup>** ± 9.9*10<sup>-5</sup><br>(2.9*10<sup>-4</sup> – 7.7*10<sup>-4</sup>) | **9.5\*10<sup>-3</sup>** ± 2.9*10<sup>-2</sup><br>(1 *10<sup>-3</sup> – 1.5*10<sup>-1</sup>) | 23.5% | 4.8% | 19.9% | 2 (18.2%) |
| Brain, frontal<br>white matter<br>($n=8$) | **0.087** ± 0.106<br>(0 – 0.377)* | **5.2\*10<sup>-4</sup>** ± 1.3*10<sup>-4</sup><br>(2.9*10<sup>-4</sup> – 7.8*10<sup>-4</sup>) | **7.3\*10<sup>-3</sup>** ± 2.8*10<sup>-2</sup><br>(1 *10<sup>-3</sup> – 1.5*10<sup>-1</sup>) | 33.6% | 12.8% | 27.9% | 5 (62.5%) |
| Cerebellum<br>($n=9$) | **0.135** ± 0.122<br>(0 – 0.419)* | **4.6\*10<sup>-4</sup>** ± 1.1*10<sup>-4</sup><br>(2.3*10<sup>-4</sup> – 6.9*10<sup>-4</sup>) | **9.4\*10<sup>-3</sup>** ± 3.0*10<sup>-2</sup><br>(1 *10<sup>-3</sup> – 1.5*10<sup>-1</sup>) | 33.5% | 15.2% | 28.2% | 3 (33.3%) |
| Placenta<br>($n=16$) | **0.28** ± 0.105<br>(0 – 0.479)* | **5.4\*10<sup>-4</sup>** ± 8.2*10<sup>-5</sup><br>(2.9*10<sup>-4</sup> – 6.8*10<sup>-4</sup>) | **4.7\*10<sup>-3</sup>** ± 3.8*10<sup>-3</sup><br>(2.5 *10<sup>-3</sup> – 2.3*10<sup>-2</sup>) | 16.9% | 12.2% | 17% | 1 (6.3%) |
| Liver<br>($n=8$) | **0.346** ± 0.101<br>(0.149 –<br>0.479) | **3.7\*10<sup>-4</sup>** ± 8.7*10<sup>-5</sup><br>(2.3*10<sup>-4</sup> – 5.9*10<sup>-4</sup>) | **2.6\*10<sup>-3</sup>** ± 2.4*10<sup>-3</sup><br>(0 – 9.9*10<sup>-3</sup>) | 14.4% | 13.8% | 16.8% | 0 (0%) |
| Lung<br>($n=9$) | **0.33** ± 0.112<br>(0.108 –<br>0.548) | **5.5\*10<sup>-4</sup>** ± 9.2*10<sup>-5</sup><br>(4 *10<sup>-4</sup> – 8.1*10<sup>-4</sup>) | **2.5\*10<sup>-3</sup>** ± 1.3*10<sup>-3</sup><br>(1 *10<sup>-3</sup> – 5.5*10<sup>-1</sup>) | 20.4% | 14.1% | 25.3% | 0 (0%) |
| Kidneys<br>($n=5$) | **0.153** ± 0.09<br>(0 – 0.307)* | **4.2\*10<sup>-4</sup>** ± 8.6*10<sup>-5</sup><br>(3.1*10<sup>-4</sup> – 6.4*10<sup>-4</sup>) | **1.7\*10<sup>-2</sup>** ± 4.2*10<sup>-2</sup><br>(1 *10<sup>-3</sup> – 1.5*10<sup>-1</sup>) | 36.2% | 17% | 30.1% | 2 (40%) |

\*: in these cases, the $n$ for assessing the variability of $f$ was further reduced due to the insufficient fitting of the bi-exponential curve. In such cases, $f$ equals zero. The number of missing data entries is given in the last column of the table.

### Factors influencing the reproducibility of IVIM parameters

In our experiments, the number of frames removed was indicative of the subject motion, and a positive correlation was found between the within-subject variability of two IVIM parameters and the number of frames removed. The number of image frames removed also influenced the reproducibility of the diffusion coefficient of the frontal cortex (ANOVA, F=6.28, p=0.046, β=0.0032) and that of the perfusion fraction of the cerebellum (ANOVA, F=25.68, p=0.007, β=0.026). Fetuses at later stages of gestation showed higher within-subject variability of the perfusion fraction of the cerebellum (ANOVA, F=14.625, p=0.019, β=0.0042). Scanner type (field strength: 1.5T or 3.0T) and maternal age were not found to correlate with any of the reproducibility measurements derived in the study.

## DISCUSSIoN

Within-subject, repeated *in utero* IVIM from 15 pregnancies demonstrated that the perfusion fraction *(f)*, diffusion coefficient *(d)* and pseudo-diffusion coefficient *(D*)* can be measured reproducibly in the fetal lungs, liver and placenta. For these organs, within-subject variability during test-retest imaging ranged from 14.4% to 20.4% for *f*, 12.2% to 14.1% for *d* and 16.8% to 25.3% for *D*.* The diffusion coefficients of the investigated brain regions were moderately to highly reproducible (4.8% to 15.2%), however, *f* and *D\** showed inferior reproducibility compared to corresponding measures derived for the lungs, liver and placenta. The IVIM-based parameters of the fetal kidney were revealed to be highly variable across scans.

### Quality of fetal IVIM data; reproducibility and confounds

The adaptation of diffusion MRI techniques to the fetal age faces numerous challenges (Kasprian et al., 2010). The IVIM approach is based on an echo-planar MRI sequence, and hence is susceptible to imaging artifacts compared to standard anatomical MRI (Le Bihan et al., 2006). Our study adds to the body of previous reports evaluating the reproducibility of the IVIM derived parameters (Grech-Sollars et al., 2015), and extends them by providing initial results *in utero.* An important part of our analysis tested whether acquisition- or subject-related confounding factors cause significant variability in the measured parameters of tissue diffusivity and perfusion. The slow diffusion coefficient *d* - the parameter that is most commonly referred to as ADC in clinical imaging studies - was the most reproducible measurement derived in our study (VAR%, placenta: 4.8, liver: 13.8%); this observation is in good agreement with previous findings from abdominal IVIM studies reporting that *d* is twice as reproducible as the microvascular perfusion fraction *f* or the *D\** (Barbieri et al., 2016). This is partially due to the fact that the mono-exponential component is estimated using numerous measurement points with higher, well separated diffusion weightings, e.g. for the present study 10 measurement points were used for high b-values (**Figure 1/b**). B-values higher than 100 s/mm^2^ are less prone to the effects of T2-relaxation. The pseudo diffusion coefficient *D\** and microvascular perfusion fraction *f* were twice as variable as *d*, highlighting the uncertainty in estimating the faster diffusion component, as in previously reported data (Pekar et al., 1992). The lower signal-to-noise ratio in estimating the fast diffusion component reduces the diagnostic value of such parameters, and a further confound arises from the fact that it may not be practically possible to reach the desired low diffusion-weighting values because of limitations of the scanner hardware (Le Bihan et al., 2006). Regarding quantitative reproducibility, considerable differences were found between fetal organs, with the brain (frontal cortex, white matter, and cerebellum) and the kidney having insufficient reproducibility for further analysis. The fetal kidneys were exceedingly prone to motion-related artifacts and their small size putatively increased partial volume related measurement errors.

The brain in the developing fetus appeared to have very low *D\** and *f* values, which can reflect low microvascular perfusion. In the adult brain, the capillary blood volume fraction is known to be very low (2-4%) compared to that in other organs (Federau et al., 2012; Le Bihan, 2012), and hence there is a need for a very good signal to noise ratio to reproducibly quantify the *D\** and *f.* Furthermore, the immaturity of cerebral capillary vasculature network in the fetus may contribute to the observed low *D\** and *f* values and their low reproducibility: in mid-gestation, the developing cortex and subcortex show lower vessel density, and vasculature is more dominated by penetrating arteries running orthogonal to the pial surface rather than long-range cortico-cortical vessels (Miyawaki et al., 1998). This may breach one of the important assumptions of IVIM imaging, namely the presence of randomly oriented vascular segments within the imaged voxels, and results in low IVIM signal in the fetal brain. However, the most likely explanation for the brain’s low (or zero) IVIM values lies in the poor data quality of these measurements. For the brain, placenta and kidney, *f* was estimated to be zero due to the insufficient fitting of the bi-exponential function in 6.3% (placenta) to 62.5% (frontal white matter) of the cases.

### Study limitations

We identified a number of additional limiting factors affecting the data quality. The availability of data for the reproducibility tests was restricted by the limited visibility of some of the organs due to the selective placement of the imaging window. Liver and lung microvascular perfusion fractions were estimated to be larger on 3.0T compared to 1.5T, as previously reported by Lemke et al. (Lemke et al., 2010) and (Park et al., 2016a; Rosenkrantz et al., 2011), allowing us to conclude that scanner field strength causes significant variability in the IVIM parameters. By the same token, the microvascular perfusion fractions were reported to be echo time dependent (Lemke et al., 2010), since longer echo times cause greater signal decay at low b-values. 3.0T is associated with larger magnetic field inhomogeneity and more susceptibility artifacts, which are exaggerated by the complex chemical environment of the amniotic fluid, the maternal organs and skeletal structures. Multi-centric studies have revealed a larger inter-scanner variability than intrascanner (for example, the IVIM parameterf had a higher intrascanner CV of 8.4% and inter-scanner CV of 24.8% in a study on brain IVIM (Grech-Sollars et al., 2015)), which calls for a careful interpretation of studies conducted on different scanners, and necessitates adequate statistical control for the possible effect of scanner type. To overcome this limitation, ROI-level or organ based estimation of IVIM parameters should be used instead of voxel level curve fitting. It was also shown that the quality of IVIM based parameters is greatly observer-dependent, and additionally depends on sequence parameters and scanner field strength (Dyvorne et al., 2013), as well as the mathematical model that is used to estimate the parameters (Conklin et al., 2016; Merisaari et al., 2017; Park et al., 2016b). In the *in utero* setting, the usability is further limited by considerable data drop-out. The need to visually control each and every image frame for motion artifacts before image processing limits the applicability of the method for diagnostic purposes. This step during the processing work-flow would optimally be replaced with the automatic evaluation of frame-to-frame motion based on similarity metrics, but such metrics are challenging to implement in practice due to the gradually changing image contrast with increasing b-values during the acquisition scheme.

**The main findings and recommendations based on our pilot, exploratory study are the following.**

Fetal IVIM measurements are moderately reproducible for the placenta, fetal lungs and liver, while the IVIM measurements were found to vary greatly in the fetal brain and kidneys, with occasional inability to fit the bi-exponential function *(f* and *D*\*=0).

The low reproducibility of the fetal brain’s IVIM parameters most likely originates relative low tissue perfusion compared to the placenta, lungs or liver.

Fetal IVIM imaging requires the optimization of the DWI sequence so that it allows for the fast acquisition of the entire fetal body and/or the placenta. In our experience, imaging times ranging from 1:40 to 3:20 are achievable.

Fetal IVIM imaging requires image processing tools that have been adapted to tackle the frame-by-frame relative movements of fetal organs due to fetal movements or maternal breathing movements. Possible solutions are nonlinear deformations or object-tracking by machine learning approaches.

An optimal strategy is to acquire diffusion-weighted images with as many b-factors as possible, preferably with NEX=1, repeated multiple times (for example, by using a tetrahedral diffusion-weighting direction scheme), and use image processing to filter out the corrupted image frames individually.

The optimal estimation of the IVIM phenomenon requires additional diffusion-weighted images with b-factors in the range of b<250 s/mm^2^.

## Acknowledgements

The authors thank the radiographer team of the Center for MR-Research for their technical assistance during the project. A.J. is supported by the OPO Foundation, the Foundation for Research in Science and the Humanities at the University of Zurich, the Hasler Foundation and the Forschungszentrum für das Kind Grant (FZK). A.J. and R.T. are supported by the EMDO Foundation, grant number 928.

